# Improved protein structure prediction by deep learning irrespective of co-evolution information

**DOI:** 10.1101/2020.10.12.336859

**Authors:** Jinbo Xu, Matthew Mcpartlon, Jin Li

**Affiliations:** Toyota Technological Institute at Chicago; Department of Computer Science, University of Chicago; University of Chicago

## Abstract

We describe our latest study of the deep convolutional residual neural networks (ResNet) for protein structure prediction, including deeper and wider ResNets, the efficacy of different input features, and improved 3D model building methods. Our ResNet can predict correct folds (TMscore>0.5) for 26 out of 32 CASP13 FM (template-free-modeling) targets and L/5 long-range contacts for these targets with precision over 80%, a significant improvement over the CASP13 results. Although co-evolution analysis plays an important role in the most successful structure prediction methods, we show that when co-evolution is not used, our ResNet can still predict correct folds for 18 of the 32 CASP13 FM targets including several large ones. This marks a significant improvement over the top co-evolution-based, non-deep learning methods at CASP13, and other non-coevolution-based deep learning models, such as the popular recurrent geometric network (RGN). With only primary sequence, our ResNet can also predict correct folds for all 21 human-designed proteins we tested. In contrast, RGN predicts correct folds for only 3 human-designed proteins and zero CASP13 FM target. In addition, we find that ResNet may fare better for the human-designed proteins when trained without co-evolution information than with co-evolution. These results suggest that ResNet does not simply denoise co-evolution signals, but instead is able to learn important sequence-structure relationship from experimental structures. This has important implications on protein design and engineering especially when evolutionary information is not available.

**Availability:** http://raptorx.uchicago.edu/ and https://github.com/j3xugit/RaptorX-3DModeling/

## Introduction

In the past few years, computational structure prediction for proteins without detectable homology in the Protein Data Bank (PDB) has been greatly advanced by the application of co-evolution analysis[1] and deep convolutional residual networks (ResNet) to facilitate accurate inter-residue contact and distance prediction[2, 3]. Since our proposal of the deep ResNet method for protein contact/distance and tertiary structure prediction[4–9], many research groups have studied this method and contributed further improvements. At CASP13, AlphaFold showed that more accurate 3D modeling can be achieved by using a deeper ResNet[10] and performing gradient-based energy minimization on predicted distance potential. Gong group showed that ResNet (coupled with GAN) can predict real-valued distance well[11]. Several groups have also capitalized on ResNet’s inherent ability to predict arbitrary pairwise relationships. Yang et al showed that deep ResNet can be used to predict inter-residue orientation[12], which may benefit 3D model building. They also show that deep ResNet works well for human-designed proteins. Moreover, Jones group showed that deep ResNet can accurately predict hydrogen-bonds, which aids in 3D structure modeling[13].

Apart from the study of model capabilities and architecture, new ideas on input features and model training have also been suggested. For example, metagenome data has been used to enhance MSAs for those proteins with very few sequence homologs[14]. A full precision matrix generated by co-evolution analysis is reported to provide more information than a concise co-evolution matrix[15]. In addition, MSA subsampling[10, 12], loop modification[16] and perturbation of distance matrices[10, 11] has also been proposed to improve model training.

Although the above-mentioned ideas have yielded promising results, we were able to obtain top performance in CASP13 for both contact prediction and 3D structure modeling with a relatively shallow ResNet, and without employing any of these ideas. In light of this, it will be interesting to see how much further the ResNet method can be improved by integrating these ideas. It will also be interesting to analyze the contribution of individual factors to further our understanding of deep learning-based protein structure prediction protocols.

Although ResNet has revolutionized structure prediction, its success is often attributed to its utilization of co-evolution information -- especially the information produced by direct-coupling analysis methods such as Evfold[17], GREMLIN[18] and CCMpred[19]. It is even suggested that the core functionality of deep ResNets is in enhancing the co-evolutionary signal derived from MSAs (multiple sequence alignment)[20]. There is little doubt that co-evolution analysis plays an important role in most successful structure prediction methods, but empirically, co-evolution analysis alone usually fails on proteins without a very deep MSA, e.g., CASP FM targets. This presents a challenge in protein design and engineering, where it is often the case that co-evolution (and even evolutionary) information is not available for engineered proteins. On the other hand, since proteins fold without knowledge of sequence homologs in nature, and thus, a method that can fold a protein in the absence of co-evolution information should exist in principle.

These considerations motivate us to study the role of co-evolution analysis with regard to deep learning in protein structure prediction. Specifically, we address how deep ResNets can be used to fold both natural and human-designed proteins with and without co-evolution information. In the case where full information is provided, we obtain significant improvements over our own CASP13 results (as well as the best CASP13 results) in both contact prediction and 3D structure modeling. Even compared to the very recent trRosetta[12], our work still yields a measurable improvement.

In the case where co-evolution information is omitted, our deep ResNet still predict correct folds for 18 of the 32 CASP13 FM targets. This marks a substantial improvement over the top co-evolution based, non-deep-learning method in CASP13 as well as the popular recurrent geometric network[20] (RGN) model which also ignores co-evolution information. The contrast is even more pronounced for human-designed proteins where our ResNet model without co-evolution may outperform its co-evolution aware counterpart (when co-evolution information is available). In fact, our deep ResNet performs well for human-designed proteins even when only primary sequence information is available. Viewed holistically, our results imply that deep ResNets do not simply denoise co-evolution signals, but instead capture fundamental aspects of protein structure essential to deciding their folded state.

### Overview of the method

Our 3D structure prediction method consists of two key components. The first is a deep ResNet used to predict discrete probability distributions over pairwise distance and inter-residue orientation. Our ResNet predicts distance distributions for three backbone atom pairs (C_b_-C_b_, C_a_-C_a_ and N-O), and three types of pairwise inter-residue orientation presented in trRosetta[12]. The second component is a revised gradient-based energy minimization method that builds 3D models from potentials derived from predicted pairwise distances, inter-residue orientations, and backbone torsion angles. Important for our study, our method is implemented so that it is easy to turn on and off specific input features and to specify which pairwise relationships should be predicted.

For each atom pair, we discretize distance into 47 bins: 0-2Å, 2-2.4Å, 2.4-2.8Å, 2.8-3.2Å,…, 19.6-20Å, >20Å, and also use one label to indicate an “unknown” distance when at least one of the two atoms do not have valid 3D coordinates in the PDB file. For orientation, we follow the same format as presented in trRosetta[12] and discretize each angle uniformly with the bin width set to 10 degrees. In addition, we use a separate orientation label for each pair to indicate that two residues are far away from each other (when the distance of their C_b_ atoms >20Å) regardless of their orientation.

The overall network architecture is similar to what we have used in CASP13[7] except that the ResNet we use here is larger and wider. Our current ResNet has 100 2D convolutional layers, each of which utilizes 150 filters on average. In comparison, our deep ResNet used in CASP13 had only 60 2D convolution layers, averaging 80 filters per layer. We simultaneously predict all distance and orientation matrices in a single deep ResNet. This multi-task learning strategy was implemented during CASP13, but not fully tested at the time. From our experience, multi-task learning does not yield a clear performance gain, but it greatly reduces the time for model testing/training as well as the number of deep model files. For all of our deep models, we used the same set of training proteins and only varied the input features (e.g., co-evolution and evolutionary information).

To build 3D models, we convert predicted distance and orientation distributions into discrete energy potential and employ an enhanced gradient descent method to minimize the sum of these potentials in the 3D model. After this, we use the fast relaxation protocol in PyRosetta[21] for side-chain packing and reducing steric clashes. We found that, on average, fast relaxation improves C_a_ conformation by around 0.01 TMscore units.

#### Accuracy of predicted contacts on CASP13 FM and FM/TBM targets

When co-evolution is used, we have trained two sets of 4 and 6 deep ResNet models using the 2018 and 2020 Cath S35 data, respectively (see data description in Methods). We use each set of deep ResNet models as an ensemble to do prediction. As shown in Table 1, in terms of both precision and F1 value of long-range contact prediction, our method outperforms the three top methods in CASP13 for FM and FM/TBM targets. This improvement is especially notable for top L/2 and top L long-range contacts, where L is the target sequence length. Our work also compares favorably to trRosetta[12], a ResNet-based program developed by Baker group after CASP13. Table 1 also shows that there is very little performance difference between the deep ResNet models trained on two different versions of Cath S35 data, which is not surprising since they are de facto not very different.

**Table 1.** Precision and F1 (%) of long-range contact prediction by several competing methods. The F1 value of trRosetta is not reported by its authors. The F1 of AlphaFold is taken from [6]. “2018” and “2020” indicate that the results are produced by our deep ResNet models trained on 2018 and 2020 Cath S35 data, respectively.

|  | 31 CASP13 FM targets |  |  | 12 CASP13 FM/TBM targets |  |  |
| --- | --- | --- | --- | --- | --- | --- |
|  | Top L/5 | Top L/2 | Top L | Top L/5 | Top L/2 | Top L |
|  | F1 of long-range contact prediction |  |  |  |  |  |
| AlphaFold in CASP13 | 22.7 | 36.9 | 41.9 | 31.4 | 48.7 | 55.1 |
| RaptorX in CASP13 | 23.3 | 36.2 | 41.1 | 28.8 | 43.2 | 51.8 |
| Zhang in CASP13 | 21.2 | 34.1 | 39.2 | 28.4 | 43.3 | 49.5 |
| This work (2020) | 27.7 | 45.1 | 52.1 | 31.3 | 52.7 | 62.5 |
| This work (2018) | 27.8 | 44.6 | 51.3 | 31.8 | 51.3 | 61.1 |
|  | Precision of long-range contact prediction |  |  |  |  |  |
| RaptorX in CASP13 | 70.0 | 58.0 | 45.0 | 85.8 | 70.1 | 56.9 |
| Zhang in CASP13 | 65.7 | 54.8 | 39.1 | 82.3 | 70.0 | 54.8 |
| trRosetta | 78.5 | 66.9 | 51.9 | NA | NA | NA |
| This work (2020) | 80.8 | 69.0 | 58.1 | 93.2 | 82.8 | 71.3 |
| This work (2018) | 80.7 | 68.7 | 57.5 | 93.4 | 81.0 | 69.6 |

##### Ablation studies

To evaluate the contribution of different factors to our method, we trained a variety of deep ResNet models using the same training set and the same procedure over different input features and two different network sizes. To save computing time, under each setting, a set of 3 deep ResNet models are trained on the 2020 Cath S35 data and then used as an ensemble to do prediction. Table 2 shows their performance on long-range contact prediction of CASP13 targets. In this table, “Large” denotes a ResNet with 100 2D convolutional layers, each layer utilizing 150 filters on average. “Small” denotes a ResNet with 60 2D convolutional layers, each of which has roughly 80 filters. “All” means that all input features are used. “No coevolution” means that neither CCMpred[19] output nor mutual information is used. “No CCMpred” means that mutual information is used, but not the CCMpred output. “No full CCMpred matrix” means that the L×L co-evolution matrix produced by CCMpred is used, but not the L×L×21×21 full precision matrix produced by CCMpred. In CASP13, we used a small ResNet, mutual information and the L×L co-evolution matrix produced by CCMpred, but not the full CCMpred matrix and the metagenome data

**Table 2.**
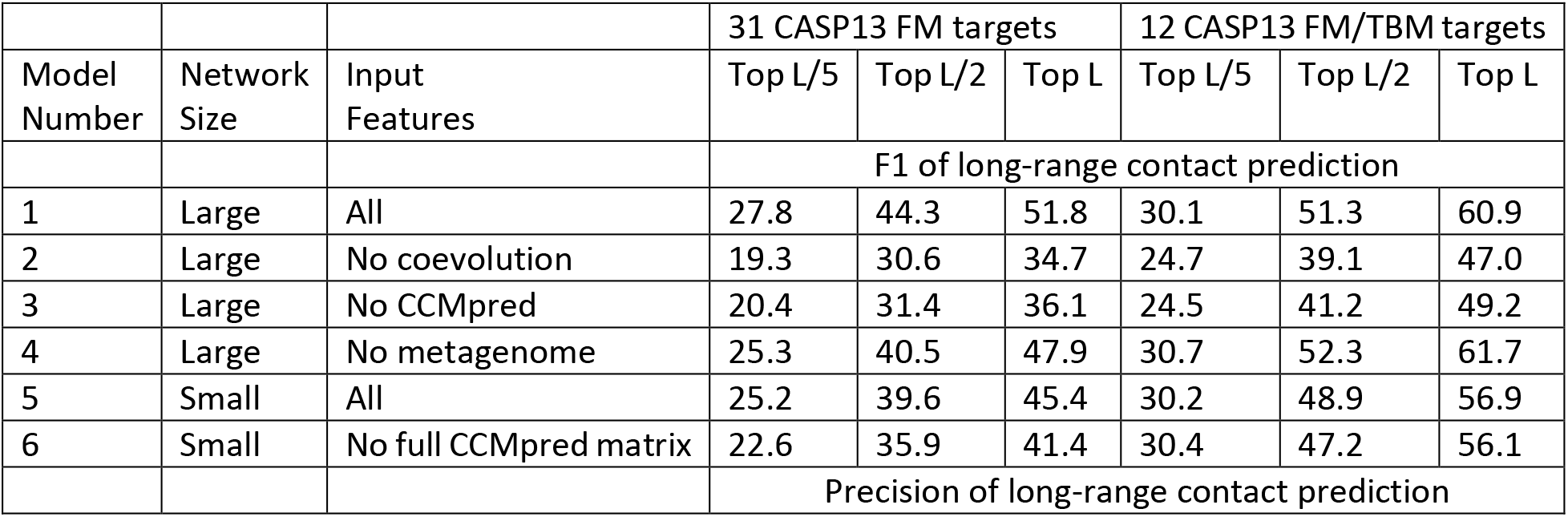
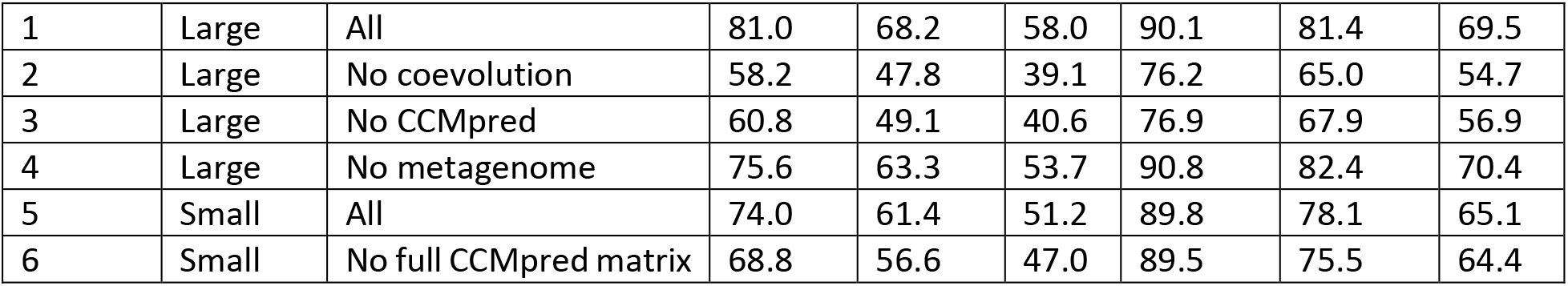
Precision and F1 (%) of long-range contact prediction on CASP13 targets by ResNet in different settings.

|  |  |  | 31 CASP13 FM targets |  |  | 12 CASP13 FM/TBM targets |  |  |
| --- | --- | --- | --- | --- | --- | --- | --- | --- |
| Model Number | Network Size | Input Features | Top L/5 | Top L/2 | Top L | Top L/5 | Top L/2 | Top L |
|  |  |  | F1 of long-range contact prediction |  |  |  |  |  |
| 1 | Large | All | 27.8 | 44.3 | 51.8 | 30.1 | 51.3 | 60.9 |
| 2 | Large | No coevolution | 19.3 | 30.6 | 34.7 | 24.7 | 39.1 | 47.0 |
| 3 | Large | No CCMpred | 20.4 | 31.4 | 36.1 | 24.5 | 41.2 | 49.2 |
| 4 | Large | No metagenome | 25.3 | 40.5 | 47.9 | 30.7 | 52.3 | 61.7 |
| 5 | Small | All | 25.2 | 39.6 | 45.4 | 30.2 | 48.9 | 56.9 |
| 6 | Small | No full CCMpred matrix | 22.6 | 35.9 | 41.4 | 30.4 | 47.2 | 56.1 |
|  |  |  | Precision of long-range contact prediction |  |  |  |  |  |
| 1 | Large | All | 81.0 | 68.2 | 58.0 | 90.1 | 81.4 | 69.5 |
| 2 | Large | No coevolution | 58.2 | 47.8 | 39.1 | 76.2 | 65.0 | 54.7 |
| 3 | Large | No CCMpred | 60.8 | 49.1 | 40.6 | 76.9 | 67.9 | 56.9 |
| 4 | Large | No metagenome | 75.6 | 63.3 | 53.7 | 90.8 | 82.4 | 70.4 |
| 5 | Small | All | 74.0 | 61.4 | 51.2 | 89.8 | 78.1 | 65.1 |
| 6 | Small | No full CCMpred matrix | 68.8 | 56.6 | 47.0 | 89.5 | 75.5 | 64.4 |

We estimate the contribution of a single factor by comparing the performance of two ResNet models in Table 2. For example, by comparing models 1 and 2, we estimate that without co-evolution information, the F1 value on average decreases by 13.1 when CASP13 FM targets are evaluated. The minor difference between models 2 and 3 implies that mutual information (one type of co-evolution information) does not help much and thus, the CCMpred output (the other type of co-evolution information) is very important. The difference between models 1 and 5 suggests that on average a large ResNet may improve F1 and precision over a small ResNet by ~4.6% and ~6.9%, respectively, on the CASP13 FM targets. By comparing models 5 and 6, we estimate that a full CCMpred matrix may improve F1 and precision by ~3.4% and ~4.7% on average, respectively. The difference between models 1 and 4 suggests that on average the metagenome data improves F1 and precision by ~3.4% and ~5.2% on the FM targets, but slightly decreases the performance on the FM/TBM targets. In summary, the factor that contributes the most to the improvement over our own CASP13 work is the network size, followed by the full CCMpred precision matrix and finally, metagenome data. We remark that these are only estimates, which may vary with respect to network size or introduction of extra features.

It is worth noting that, when co-evolution is not used, the top L/5, L/2 and L long-range contact precision obtained by our ResNet on the 31 CASP13 FM targets is 58.2%, 47.8% and 39.1%, respectively. Compared to in CASP11, where traditional neural networks utilizing coevolution (via direct-coupling analysis), such as MetaPSICOV[22], and deep belief networks (e.g., DNCON[23]) were employed, we significantly outperform the best reported precision, i.e., ~27% for top L/5. In addition, we outperform the best top L/5 accuracy (~47%) in CASP12 where an under-developed co-evolution aware deep ResNet model was tested in RaptorX-Contact server[5]. Even in CASP13 when deep ResNet with co-evolution was implemented by many top-placing groups, a method with top L/5 long-range precision of 58.2% would still be ranked among top 10. This strongly suggests that low-precision contact prediction prior to CASP12 cannot be attributed to the fact that co-evolution was not widely used.

### Accuracy of predicted 3D models on CASP13 FM targets when co-evolution is used

We test our (large) ResNet by generating 150 decoys per target and clustering them based on a pairwise RMSD cutoff. Again, here we evaluate two sets of 4 and 6 deep ResNet models trained on the 2018 and 2020 Cath S35 data, respectively. Each ResNet has 100 2D convolutional layers and each of which has 150 filters on average. When the set of 6 ResNet models trained on the 2020 Cath S35 data is used, the average quality (TMscore) of the first model (the first cluster centroid) and the best model (the best cluster centroid) is 0.629 and 0.663, respectively. When the set of 4 ResNet models trained on the 2018 Cath S35 data is used to generate 150 decoys per target, the average quality of the first and best models is 0.638 and 0.659, respectively. This result further confirms that in terms of prediction performance there is no big difference between the two Cath S35 datasets. The modeling accuracy can be further improved. For example, increasing to 600 decoys per target may improve the first and best model quality to 0.640 and 0.675, respectively. Increasing the 2D ResNet size to 120 convolutional layers (and 170 filters per layer) may improve the first and best model quality to 0.646 and 0.673 with even fewer than 150 decoys generated per target.

As shown in the left picture of Fig. 1, our predicted 3D models for 24 of 32 CASP13 FM targets have TMscore higher than their respective target-training structure similarity (i.e., the highest structure similarity between a specific target and all training/validation proteins). For 18 targets, their predicted models have quality at least 0.1 (in terms of TMscore) higher than their respective target-training structure similarity. There is almost no correlation between our model quality and the target-training structure similarity (correlation coefficient=−0.149 and trendline R^2^=0.0223) (Fig. 1, left). When the best models are considered and TMscore>0.5 is used to judge if a predicted 3D model has a correct fold or not, our method predicts correct folds for 26 of the 32 FM targets. These results demonstrate the power of ResNets in generating novel structures, yielding model quality remarkably higher than what can be achieved by simply memorizing the training set. There is a modest correlation between model quality and MSA depth (correlation coefficient=0.572 and trendline R^2^=0.3276) (Fig. 1, right) and our method predicts correct folds for all but two test targets with Meff >2, (i.e., MSA depth>7.39).

**Figure 1.**
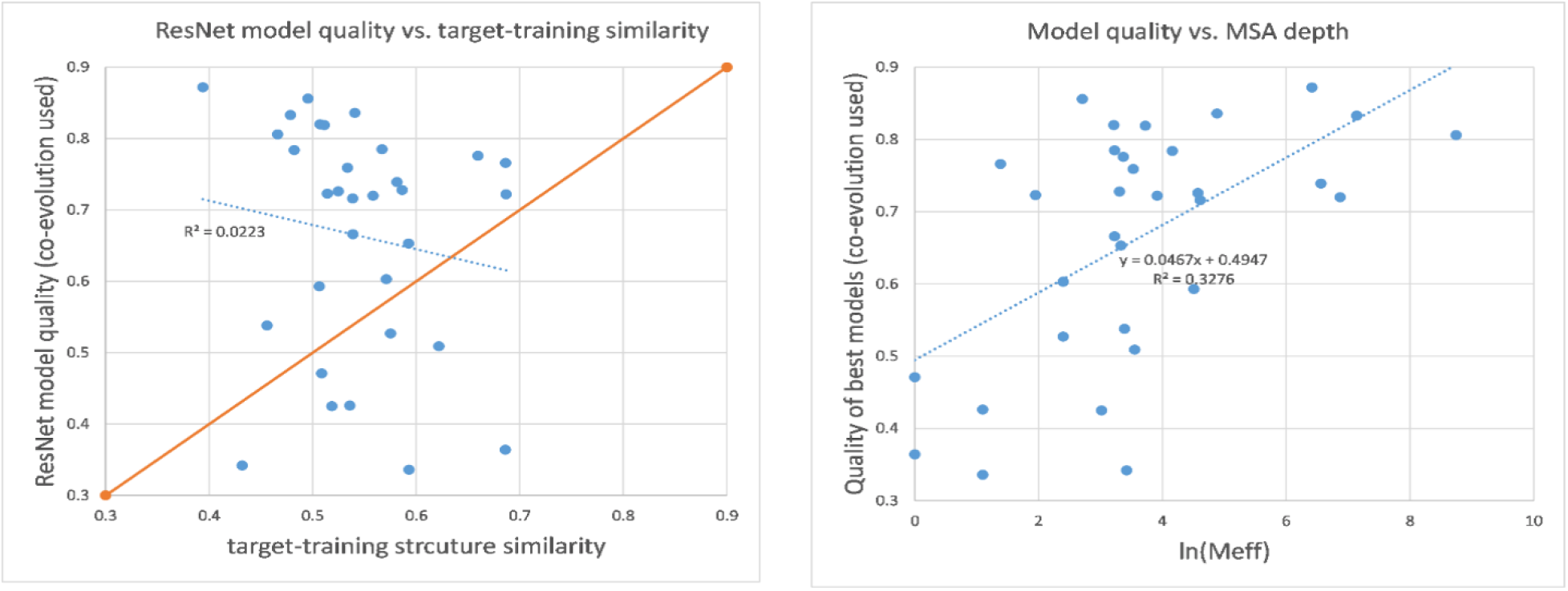
3D modeling accuracy (measured by TMscore) on CASP13 FM targets when co-evolution is used with ResNet. Left: relationship between our 3D model accuracy and target-training structure similarity. A dot above the diagonal line indicates that the quality of our predicted 3D model quality is larger than the highest target-training structure similarity. Right: relationship between our 3D model accuracy and MSA depth (i.e., Meff). The results are obtained when 150 decoys are generated per target by the set of 6 deep ResNet models trained on 2018 Cath S35 data.

Our 3D modeling accuracy compares favorably to previously reported results. For example, in CASP13 AlphaFold predicted correct folds for 23 of 32 CASP13 FM targets with average quality (TMscore) for the first and best models of 0.583 and 0.625, respectively. The lowest-energy models produced by trRosetta on the 32 CASP13 FM targets have average TMscore 0.618. Note that trRosetta reported an average quality (0.625) of 31 (but not 32) CASP13 FM targets in [12].

When the orientation potential is not used, the average quality (TMscore) of the first and best models generated by our method is 0.628 and 0.648, respectively, when the set of 6 deep ResNet models are used. That is, using orientation potential may improve the best model quality by about 0.015 TMscore unit. In contrast, trRosetta reported a larger improvement resulting from orientation potential [12]. This may be because 1) we use distance potential for three types of atom pairs whereas trRosetta used only C_b_-C_b_ distance; and 2) we make use of orientation information in a much simpler way than what is described in the trRosetta paper.

### Folding proteins without co-evolution information

Co-evolution information plays an important role in recent progress on protein structure prediction. Here we study how well a ResNet can fold a protein when co-evolution is not used. Note that although co-evolution is not used, we still make use of sequence profile (one kind of evolutionary information), unless explicitly stated otherwise. Sequence profile encodes evolutionary information on an individual residue basis while co-evolution encodes evolutionary information on two residues. We have trained 4 large ResNet models of similar size (i.e., 100 2D convolutional layers and 150 filters per layer) on the 2018 Cath S35 data using the same set of training proteins and the same procedure, but without co-evolution. We test this kind of ResNet models on the 32 CASP13 FM targets and 21 human-designed proteins.

#### Folding CASP13 FM targets without co-evolution information

The average quality (measured by TMscore) of the first and best models generated by our ResNet models is 0.478 and 0.506, respectively. When the best models are considered and a model with TMscore>0.5 is assumed to have a correct fold, our deep ResNet predicts correct folds for 18 out of 32 CASP13 FM targets (Fig. 2). There is a weak correlation between 3D modeling accuracy and target-training structure similarity (correlation coefficient=0.363 and trendline R^2^=0.1315), as shown in the left picture of Fig. 2. The figure also shows that the correlation between modeling accuracy and MSA depth is weaker (correlation coefficient=0.211 and trendline R^2^=0.0448) than when co-evolution is used (Fig. 1). That is, when co-evolution is not used, 3D modeling accuracy depends less on the number of sequence homologs, which is not surprising.

**Figure 2.**
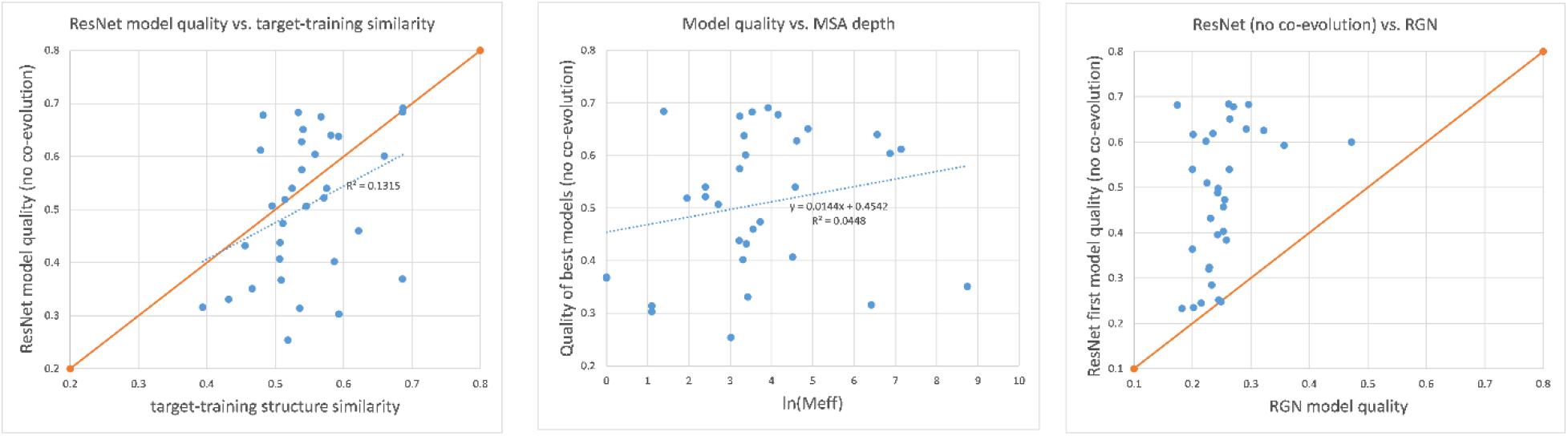
3D modeling accuracy on CASP13 FM targets when co-evolution is not used. Left: relationship between our 3D model accuracy and target-training structure similarity. A dot above the diagonal line indicates predicted 3D model quality is larger than target-training structure similarity. Middle: relationship between modeling accuracy and MSA depth (i.e., Meff). Right: comparison of our 3D modeling accuracy with RGN. A dot above the diagonal line indicates that the model predicted by ResNet without co-evolution has higher quality than its corresponding model by RGN.

Fig. 2 also shows that deep ResNet can predict structures for 13 FM targets with quality higher than their respective target-training structure similarity. In particular, the best models generated by our deep ResNet for T0950-D1 (342AAs), T0968s2-D1 (115AAs), T0969-D1 (354AAs), T0986s2-D1 (115AAs), T0987-D1 (185AAs) and T1017s2-D1 (125AAs) have TMscore 0.628, 0.651, 0.612, 0.683, 0.675, and 0.678, respectively, while their target-training structural similarity is only 0.539, 0.540, 0.478, 0.533, 0.567, and 0.482, respectively. Among the 3 FM targets with more than 300 residues (T0950-D1, T0969-D1 and T1000-D2), deep ResNet predicts correct folds for T0950-D1 and T0969-D1, both of which have predicted model quality much higher than their respective target-training structure similarity. To understand why deep ResNet works well for T0969-D1 (a very large test protein without a similar fold in our training set) when co-evolution is not used, we visualize its predicted backbone atom distance matrices in Fig. 3. Though less precise than the predicted distance matrix obtained when co-evolution is used, this figure shows that in the absence of co-evolution information, deep ResNet can still correctly predict a good percentage of long-range distances, even for pairs of residues with sequence separation exceeding 200 residues. In summary, our results indicate that even without co-evolution information, deep ResNet is still able to predict the correct fold structure for many natural proteins.

**Figure 3.**
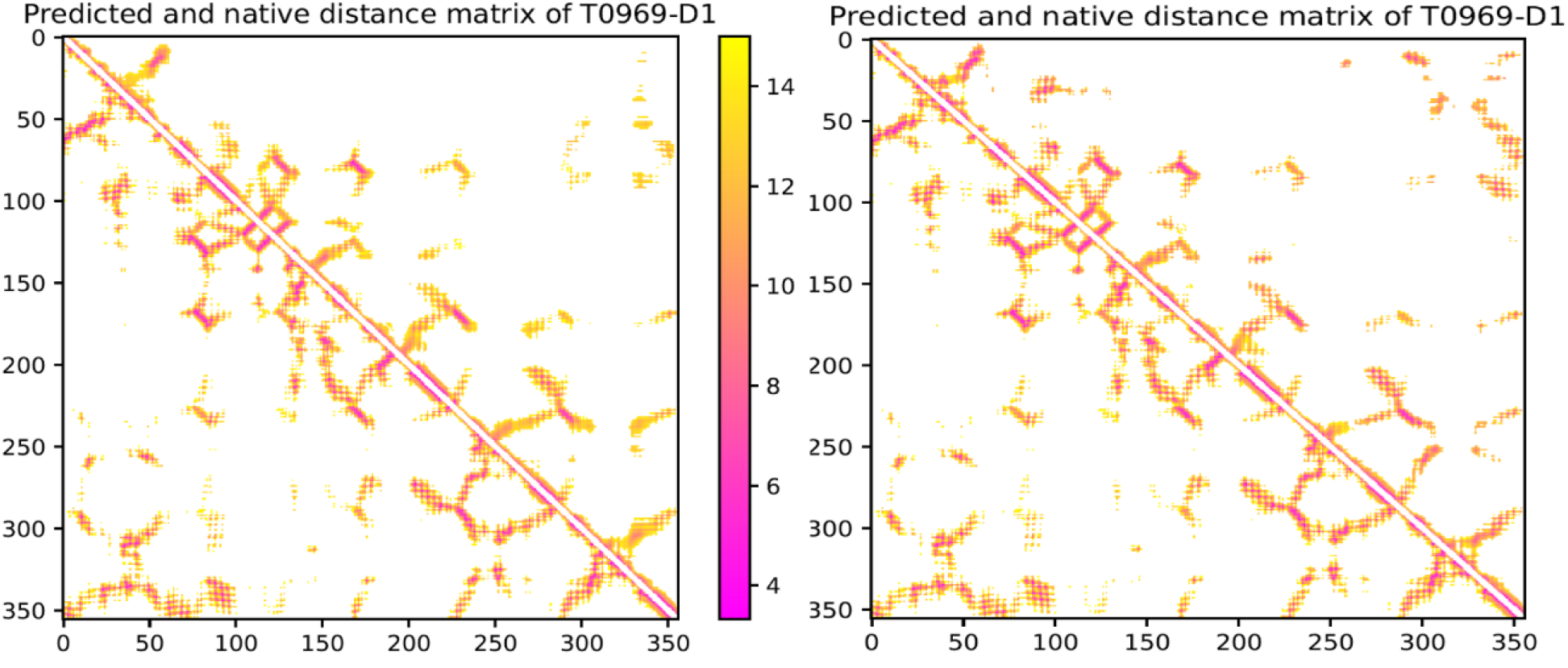
Distance matrices predicted for T0969-D1 by deep ResNet when co-evolution is not used (left) and used (right). Only distance predictions less than 15Å are displayed in color. In each picture, native distance is shown below the diagonal, and predicted distance is shown above.

To put our work into perspective, we compare it with a top server Robetta[24]in CASP13 and a popular deep learning method RGN (recurrent geometric networks) [25]. Robetta is the best server in CASP13 that did not make use of deep learning, but used a combination of template-based modeling, ab initio folding and co-evolution-based contact prediction. Both template-based modeling and ab initio folding are built upon sequence profile, i.e., evolutionary information at individual residues. RGN is a popular end-to-end deep learning method that predicts a single 3D structure of a protein from its sequence profile without using co-evolution information. In this work, the RGN results are generated by its pre-trained deep model for the CASP12 targets. We use the uniref90 database dated in November 2018 to generate sequence profile of a test protein as input of the RGN deep model.

The average quality (measured by TMscore) of the first and best models submitted by Robetta on the 32 CASP13 FM targets is 0.390 and 0.430, respectively. The average quality of the 32 RGN models is 0.251. Compared to our deep ResNet - with first and best models having average TMscore 0.478 and 0.506 - both methods produce models with significantly lower TMscore. Furthermore, our ResNet trained without co-evolution predicts 3D models with better quality than RGN for almost all 32 FM targets (Fig 2, right). In fact, RGN fails to produce a correct fold for all the 32 test targets, while Robetta and our ResNet predict 7 and 15 correct folds, respectively, when only the first models are considered. We also point out that, when only primary sequence along with its derived secondary structure and solvent accessibility is used as input to our deep ResNet (trained with sequence profile), we are only able to predict correct folds for 2 of the 32 FM targets. For this method, the average TMscore is only ~0.30 across the FM targets. That is, for natural proteins our current implementation of the deep ResNet method does not fare well when only primary sequence is available.

#### Folding human-designed proteins

Baker group has shown that deep ResNet can fold 16 of the 18 de novo proteins designed by the same group[12]. Here we conduct a more comprehensive study on 21 de novo proteins designed by two research groups in 2018-2020[26–28], among which 11 proteins have been used to test trRosetta. None of these 21 proteins has evolutionarily related homologs in the 2018 Cath S35 training proteins (HHblits with E-value<0.1). We consider two types of ResNet models. One is trained and tested with all input features including co-evolution information. The other is trained and tested without co-evolution information, mainly using input features derived from sequence profile. We also test two types of input features. One is derived from MSAs (multiple sequence alignment) built by HHblits on the 2018 uniclust30 sequence database and the other from MSAs built by HHblits on the 2020 uniref30 sequence database. In fact, no sequence homologs are found for the 21 test proteins in the 2018 database and few homologs are found for 9 test proteins in the 2020 database.

Table 3 summarizes the modeling accuracy of the two types of ResNet models with two types of input features. When evolutionary information is not available at all (i.e., the 2018 uniclust30 database is used to find sequence homologs), our ResNet still can predict correct folds (i.e., TMscore>0.5) for almost all the 21 proteins no matter whether it is trained with or without co-evolution information (Fig. 4, top left and top right). Table 3 and Fig. 4 also show that evolutionary information (if available) helps improving modeling accuracy, especially when only the first 3D models are evaluated and ResNet is not trained with co-evolution information.

**Table 2.** Average modeling accuracy of deep ResNet on 21 human-designed proteins. GDT and GHA are scaled to [0, 1].

| ResNet trained with | Seq database used to build input features | First Model |  |  | Best Model |  |  |
| --- | --- | --- | --- | --- | --- | --- | --- |
|  |  | TM | GDT | GHA | TM | GDT | GHA |
| Co-evolution | 2018 uniclust30 | 0.650 | 0.662 | 0.479 | 0.675 | 0.686 | 0.498 |
| Co-evolution | 2020 uniref30 | 0.662 | 0.675 | 0.490 | 0.683 | 0.694 | 0.509 |
| No coevolution | 2018 uniclust30 | 0.584 | 0.598 | 0.430 | 0.676 | 0.695 | 0.509 |
| No coevolution | 2020 uniref30 | 0.654 | 0.663 | 0.495 | 0.723 | 0.738 | 0.556 |

**Figure 4.**
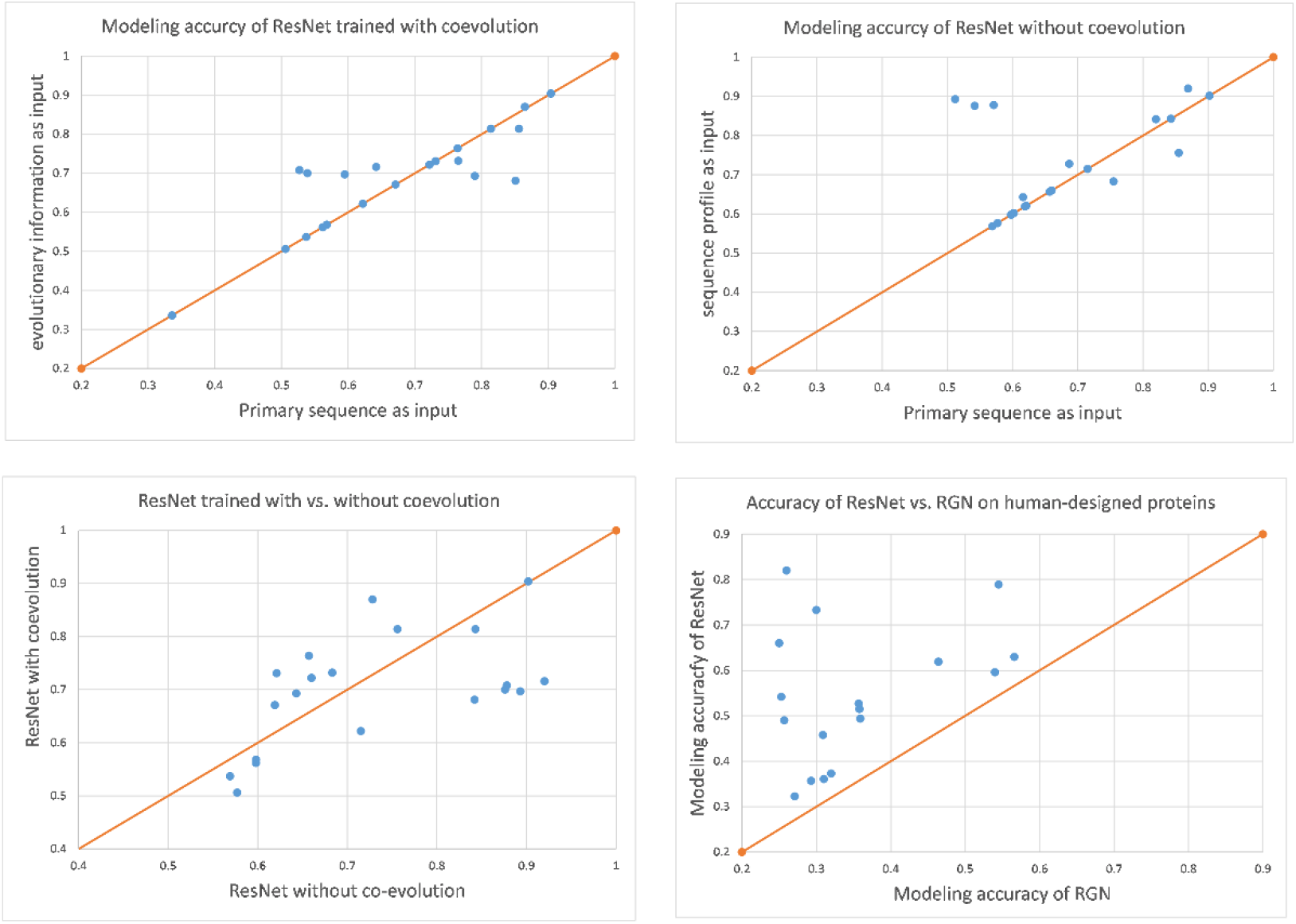
3D modeling accuracy on the human-designed proteins. Top Left: modeling accuracy by deep ResNet trained with co-evolution. A dot above the diagonal line indicates that a 3D model predicted with evolutionary information (coevolution and sequence profile) has higher quality. Top Right: modeling accuracy of deep ResNet trained without co-evolution. A dot above the diagonal line indicates that a 3D model predicted with evolutionary information (coevolution and sequence profile) has higher quality. Bottom Left: modeling accuracy of ResNet trained with vs. without co-evolution. A dot above the diagonal line indicates that a 3D model predicted by ResNet trained with co-evolution has higher quality. Bottom Right: ResNet vs. RGN. ResNet is not trained with co-evolution information and primary sequence is used as input. A dot above the diagonal line indicates that the first model predicted by ResNet for a specific target has higher quality.

When very few sequence homologs are available (i.e., the 2020 uniref30 database is used to find sequence homologs) and the best 3D models are considered, ResNet trained without co-evolution outperforms ResNet trained with co-evolution (Fig. 4, bottom left). The average TMscore of the best 3D models obtained by ResNet without co-evolution and with co-evolution is 0.723 and 0.683, respectively. This may be because that when very few sequence homologs are available, co-evolution information is not as reliable as sequence profile.

The 21 models generated by RGN (recurrent geometric networks) have average TMscore 0.363, much worse than the average quality of the first models predicted by ResNet trained without co-evolution and using primary sequence as input (Fig. 4, bottom right). In total RGN predicts correct folds for only 3 of the 21 human-designed proteins while ResNet predicts 14 correct folds when the first models are considered. In this work, the RGN results are generated by its pre-trained deep model for the CASP12 targets. We use the uniref90 database dated in November 2018 to generate sequence profile of a test protein as input of the RGN deep model.

Yang et al tested trRosetta on 11 of the 21 human-designed proteins. On these 11 proteins, trRosetta predicted 3D models with average TMscore 0.661, comparable to our ResNet trained with co-evolution, which is not very surprising since trRosetta also uses a ResNet trained with co-evolution information.

## Conclusion and Discussions

We have presented our latest study of deep ResNet for protein contact/distance prediction and template-free protein folding. Our experimental results show that, although in the past several years protein contact and tertiary structure prediction has been significantly improved by deep ResNet, further improvements can still be made by integrating a variety of ideas. On the CASP13 FM targets, we obtain top L/5 long-range contact precision over 80%, improving over the best CASP13 results by roughly 10%. Additionally, we can predict correct folds for all but two CASP13 FM targets with at least 7 sequence homologs (nonredundant at 70% sequence identity level). Compared to our work in CASP13, the major improvements in contact prediction results from using a larger ResNet, training on the full precision matrix generated by direct coupling analysis, and by using metagenome data. Although no single idea improved the performance by a very large margin, the combination of these ideas results in measurable improvement. The improvement in 3D structure modeling is likely attributed to the use of gradient-based energy minimization and the incorporation of inter-residue orientation information. Our 3D modeling procedure is still very simple, less sophisticated than what is described in[12], especially in the application of predicted inter-residue orientation information. In future work, we will investigate the full effect of orientation information for 3D modeling.

We have shown that when co-evolution is not used, a deep ResNet trained with sequence profile can predict correct folds for more than half of the CASP13 FM targets and all the human-designed proteins tested by us. Further, we found that, for the human-designed proteins, sequence profile is more useful than co-evolution information when few sequence homologs are available. In the case where evolutionary information is not available (i.e., only primary sequence and its derived properties is used as input), deep ResNet still can predict correct folds for almost all the human-designed proteins, albeit with a loss in average model quality. These results clearly indicate that the deep ResNet model learns important information about protein folding from experimental protein structures and the functionality of ResNet extends beyond denoising co-evolution signals.

Although our deep ResNet can predict correct folds for most of the human-designed proteins without using any evolutionary information, it does not work well on natural proteins. This may be because that the folds of human-designed proteins are often well optimized by human experts, whereas natural proteins are not. For most natural proteins, our method still requires a small number of sequence homologs to obtain correct fold prediction. Nevertheless, in nature a protein folds without knowledge of its sequence homologs, and it is desirable to have a method that can fold a protein without resorting to using sequence homologs. Recently, language models such as Transformer have been used to model protein sequences by a few research groups[29]. It will be interesting to determine the use of this model for aiding in structure prediction when few homologs are present.

## Methods

### Training and validation data

In CASP13 we trained deep ResNet models using PDB25, a set of representative protein chains generated by the PISCES server[30] in which no two protein chains share >25% sequence identity. Here we use Cath S35 as our training and validation proteins. Cath S35 is a representative set of CATH domains (https://www.cathdb.info/), in which any two domains share no more than 35% sequence identity. According to our previous work[6], PDB25 and Cath S40 result in similar performance on template-free modeling (FM) targets, but Cath S40 may have a slightly better performance on template-free/template-based modeling (FM/TBM) targets. The protein domains in Cath S35 on average are shorter than the protein chains in PDB25, so it may reduce GPU memory consumption by using Cath S35. For each protein domain in Cath S35, we generated their multiple sequence alignments (MSAs) by running HHblits with E-value=0.001 on the uniclust30 library dated in 2017 and then derived input features for the prediction of distance and orientation.

We downloaded two versions of Cath S35 at ftp://orengoftp.biochem.ucl.ac.uk/cath/releases/daily-release/archive/. One is dated in March 2018 and the other on January 1, 2020. They have 32140 and 32511 entries, respectively, and differ by about 2000 entries. We excluded very short domains (<25 AAs) and those with too many missing C_a_ and C_b_ atoms and then randomly split them into two non-overlapping subsets: one for training and the other for validation (1800 domains). In total we generated 6 splits, but not all of them were used to train our deep learning models due to limit of computing resources. Our results show that there is little difference between these two Cath S35 datasets in terms of both contact prediction and 3D modeling (see Results section).

### Independent test data

#### Target-training similarity

We use HHblits[31] with E-value=0.1 to check if there are evolutionarily related proteins in our training/validation data for a specific test target. When the E-value returned by HHblits is larger than 0.1, we say that this target has no evolutionarily related proteins in our training set. We also use TMalign[32] to calculate target-training structure similarity, which is defined as the highest structure similarity (measured TMscore) between a specific target and all our training/validation proteins. By the way, we always use the target length (i.e., the number of valid C_a_ atoms in an experimental structure) as the normalization constant while using TMalign to calculate structure similarity and the TMscore program to evaluate the quality of one predicted 3D model, so that TMscore returned by TMalign and the TMscore program is comparable.

#### CASP13 FM and FM/TBM targets

In total there are 32 CASP13 FM targets and 13 FM/TBM targets, among which one FM target T0950 is a server-only target. T0953s1 and T0955 have very few long-range contacts, so they are not used to evaluate contact prediction. We mainly use the CASP13 FM targets to evaluate our method since some FM/TBM targets may have evolutionarily related proteins in PDB dated before CASP13. We use HHblits (with E-value=0.1) and TMalign to check sequence profile and structure similarity between the CASP13 FM targets and our training set (i.e., the two Cath S35 datasets). HHblits returns a large E-value (>10) for many of the 32 FM targets. Only two FM targets T0975 and T1015s1 have HHblits E-value less than 0.1. T0975 is related to 4ic1D with HHblits E-value=4.2E-12, and T1015s1 is related to 4iloA with HHblits E-value=0.024. Both 4ic1D and 4iloA were deposited to PDB well before 2018, so it is fair to include them in our training set. Further, the structure similarity (TMscore) between T0975 and 4ic1D and between T1015s1 and 4iloA is less than 0.5.

#### De novo proteins

We collected 35 de novo proteins (including 7 membrane proteins) designed by two research groups in the past several years. Meanwhile, 4 of them are designed by Kortemme group[28] and the others by Baker group. Among these proteins, 21 of them have HHblits E-value>0.001 with our training proteins in the 2018 Cath S35 dataset, so we only use these 21 proteins as our test set. Their PDB codes are 6B87, 6CZG, 6CZH, 6CZI, 6CZJ, 6D0T, 6DG5, 6DG6, 6E5C, 6M6Z, 6MRR, 6MRS, 6MSP, 6NUK, 6O35, 6TJ1, 6TMSA, 6TMSG, 6VG7, 6VGA and 6VGB. To further remove overlap between our training set and these test proteins, we exclude 20 proteins from the 2018 Cath S35 dataset so that all the 21 test proteins have HHblits E-value>0.1 with our training set. All our deep ResNet models used to predict these 21 de novo proteins are trained by this Cath S35 dataset. Among the 21 proteins, 11 of them have been used by Baker group to evaluate trRosetta [12]. They are 6CZG, 6CZH, 6CZI, 6CZJ, 6D0T, 6DG5, 6DG6, 6E5C, 6MRR, 6MRS, 6MSP, and 6NUK.

We run HHblits (E-value=0.001) to search through two sequence databases released by Soding group (i.e., uniclust30 dated in March 2018 and uniref30 dated in February 2020) to see if there are any sequence homologs for the 21 test proteins. It turns out that HHblits does not find any sequence homologs in the 2018 uniclust30 database possibly because these 21 proteins are released after March 2018. When the uniref30 database is used, HHblits finds few sequence homologs for 9 of the 21 proteins (6B87, 6CZG, 6CZH, 6CZI, 6CZJ, 6D0T, 6VG7, 6VGA and 6VGB). Further examination indicates that most of these sequence homologs are also human-designed proteins or themselves.

### MSA generation and input features

We run HHblits[33] with E-value=1E-3 and 1E-5 and jackhammer[34] with E-value=1E-3 and 1E-5 to detect sequence homologs and generate MSAs (multiple sequence alignment) for CASP13 FM targets. To ensure a fair comparison, we generated their MSAs using the uniclust30 database dated in October 2017 and the uniref90 database dated in March 2018. In addition, when a CASP13 target has a shallow MSA (i.e., ln(Meff)<6), we also search a metagenome database dated in May 2018 to see if more sequence homologs can be found. The metagenome data is not applied to the 21 human-designed proteins.

We use the following input features: 1) primary sequence: one-hot encoding is used to represent a sequence; 2) sequence profile: it is derived from MSA and encodes evolutionary information at each residue; we also use secondary structure and solvent accessibility predicted from sequence profile; 3) co-evolution information: mutual information and CCMpred output. Meanwhile, the CCMpred output includes one L×L co-evolution matrix and one full precision matrix of dimension L×L×21×21 where L is the protein sequence length. We have also tested one covariance matrix of dimension L×L×21×21, but not observed performance gain and thus, decided not to use it. In CASP13, we only used the L×L co-evolution matrix generated by CCMpred as an input feature, but not the full precision matrix. When sequence profile is not used, we use secondary structure and solvent accessibility predicted from primary sequence. For each CASP13 target, there are 4 different MSAs generated. We run our deep ResNet on these 4 MSAs separately to predict 4 sets of distance and orientation and the calculate their average to obtain the final prediction.

#### MSA depth

We use *Meff* to measure the depth of an MSA, i.e., the number of non-redundant (or effective) sequence homologs in the MSA. Let *S*_*ij*_ be a binary variable indicating whether two protein sequences *i* and *j* are similar. *S*_*ij*_ is equal to 1 if and only if the sequence identity between *i* and *j* is >70%. For a protein *i*, let *S*_*i*_ denote the sum of *S*_*i1*_, *S*_*i2*_,…, *S*_*i,n*_ where *n* is the number of proteins in the MSA. Then, *Meff* is calculated as the sum of *1/S*_*1*_, *1/S*_*2*_,…,*1/S*_*n*_. Generally speaking, the smaller *Meff* a target has, the more challenging to predict its contact and distance. MSA depth of a specific target depends on the homology search tool, sequence database and E-value cutoff. In this paper, for CASP13 targets we the MSAs generated by HHblits with E-value=0.001 on the uniclust30 database dated in October 2017.

### Protein structure representation

In this work we represent only protein backbone conformation using inter-atom distance matrices and inter-residue orientation matrices. Three types of Euclidean distance matrices are used for three types of atom pairs: C_a_-C_a_, C_b_-C_b_ and N-O. In addition to the inter-residue orientation defined by N, C_a_ and C_b_ atoms of two residues and implemented in trRosetta, we have studied another type of orientation formed by four C_a_ atoms (of 4 residues): C_a_(i), C_a_(i+1), C_a_(j) and C_a_(j+1). This C_a_-based orientation includes one dihedral angle formed by two planes C_a_(i)-C_a_(i+1)-C_a_(j) and C_a_(i+1)-C_a_(j)-C_a_(j+1) and two angles formed by 3 atoms, i.e., C_a_(i), C_a_(i+1), C_a_(j) and C_a_(j), C_a_(j+1), C_a_(i). The advantage of this C_a_-based orientation is that it involves 4 residues instead of only 2 residues, no special handling is needed for Glycine, and it also allows us to use those protein structures with many missing C_b_ atoms to train our deep models. However, our experimental results show that this C_a_-based orientation slightly underperforms the orientation employed by trRosetta, so by default we employ the inter-residue orientation implemented in trRosetta. We discretize distance between 2 and 20Å uniformly into 45 bins (i.e., binwidth=0.4Å) and orientation uniformly by 10 degree.

### Deep model training

Similar to AlphaFold and trRosetta, we train our deep ResNet by subsampling MSAs. Each time we randomly sample 50% of the sequence homologs from an MSA (when it has at least 2 sequences) and then derive input features from the sampled MSA including sequence profile and co-evolution information. We train our deep ResNet for 20 epochs and select the model with the minimum validation loss as our final model. To examine the impact of input features and network capacity, we have trained deep ResNet models with different input features and network sizes. Under each setting, we trained up to 6 deep ResNet models and used them as an ensemble to predict distance and orientation probability distributions.

### Building protein 3D models and model clustering

The 3D model building protocol consists of the following major steps: (1) convert predicted distance and orientation probability distribution into discrete energy potential using the method described in our previous work [35]. We have experimented with both DFIRE[36] and DOPE[37] reference states and found that on average the DFIRE reference state is slightly better. For orientation potential, we obtain its reference state by simple counting the occurring frequency of each orientation label in our training proteins. (2) interpolate the discrete energy potential for each pair of residues to a continuous curve using the Rosetta spline function; (3) minimize the energy potential by gradient-based energy minimization, i.e., the LBFGS algorithm implemented in Rosetta. The starting 3D model for energy minimization is sampled from the phi/psi distribution predicted by our deep learning method (implemented before CASP13) from sequence profile. Since LBFGS may not converge to the global minimum, once it reaches a local minimum, we perturb all phi/psi angles by a small deviation and then apply LBFGS again to see if a conformation with a lower energy potential may be generated. This perturbation procedure is repeated up to three times and may slightly improve modeling accuracy. After this LBFGS step, a fast relaxation protocol is applied to add side-chain atoms and reduce steric clashes. The energy minimization and relaxation steps are implemented with PyRosetta.

We generated 150 decoys for each test target and then used SPICKER[38] to cluster them into up to 10 groups. The first and the best cluster centroids are used as the first and best predicted 3D models, respectively. There is only very minor decrease in modeling accuracy when the best of top 5 (instead of top 10) cluster centroids is used as the best model.

### Performance metrics

Following CASP official assessment, we use precision and F1 value to evaluate the long-range contact prediction. We mainly use TMscore[39] to evaluate the quality of a 3D model, which measures the similarity between a 3D model and its experimental structure (i.e., ground truth). It ranges from 0 to 1 and usually a 3D model is assumed to have a correct fold when its TMscore≥0.5. The GDT and GHA scores of a 3D model are also presented along with TMscore, although we do not focus on them. By definition, GDT and GHA range from 0 to 100, but we normalize them by 100 to have the same scale as TMscore.

## Acknowledgements

The authors are grateful to Drs. Jianyi Yang and Ivan Anishchanka for their very helpful discussions, providing trRosetta results and helping with PyRosetta.

## Author Contributions

J.X. conceived the whole project, implemented and tested the code, and wrote the manuscript. M.M. studied the gradient-based energy minimization algorithm and revised the manuscript. J.L. studied the deep learning algorithms and generated the RGN results.

